# AirLift: A Fast and Comprehensive Technique for Remapping Alignments between Reference Genomes

**DOI:** 10.1101/2021.02.16.431517

**Authors:** Jeremie S. Kim, Can Firtina, Meryem Banu Cavlak, Damla Senol Cali, Nastaran Hajinazar, Mohammed Alser, Can Alkan, Onur Mutlu

## Abstract

AirLift is the first read remapping tool that enables users to quickly and comprehensively map a read set, that had been previously mapped to one reference genome, to another similar reference. Users can then quickly run downstream analysis of read sets for each latest reference release. Compared to the state-of-the-art method for remapping reads (i.e., full mapping), AirLift reduces the overall execution time to remap read sets between two reference genome versions by up to 27.4×. We validate our remapping results with GATK and find that AirLift provides high accuracy in identifying ground truth SNP/INDEL variants.

**Code Availability:** AirLift source code and readme describing how to reproduce our results are available at https://github.com/CMU-SAFARI/AirLift.

## 1. Introduction

Reference genomes are inaccurate and do not perfectly represent the average healthy individual of a species for a variety of reasons [1, 2]. First, reference genomes are constructed using imperfect sequencing technologies that result in error-prone reads [3]. Second, the sequenced reads of an individual (i.e., *read set*) are assembled into a reference genome using imperfect assembly tools [4, 5]. As genome sequencing technology and assembly algorithms improve, and as more sequenced samples become available, researchers are able to incrementally assemble more accurate reference genomes. As an example, the Genome Reference Consortium (GRC) releases minor updates to the human reference genome every three months and major updates every few years [6, 7]. Very recently, significant advances have resulted in a novel full telomere to telomere reference [8]. These updates are *critical* to the accuracy of the reference genome as they enable the latest reference genome to provide the most accurate and complete representation of the reference’s respective population. Therefore, a read set should be mapped to the latest and most relevant reference genome to obtain the most accurate downstream genome analysis results [9].

Currently, the best way to adapt an existing genomic study (i.e., read sets from many samples) to a new reference genome is to re-run the *entire* analysis pipeline using the new reference genome. For example, the original analysis of the read sets from the 1000 Genomes Project was completed using the human reference genome build 37 (GRCh37) [10]. After the next version of the reference (GRCh38) became available, each read set from the 1000 Genomes Project was mapped again to the new human reference genome (GRCh38) [11]. Unfortunately, this approach is *computationally very expensive* and does not scale to large genomic studies that include a large number of individuals for three key reasons. First, mapping even a *single* read set is computationally expensive [12, 13] (e.g., 75 hours for aligning 300,000,000 short reads, which provides 30*×* coverage of the human genome) as it heavily relies on a computationally-costly alignment algorithm [14, 15, 16]. Second, the number of available read sets doubles approximately every 8 months [17, 18], and the rate of growth will continue to increase as sequencing technologies continue to become more cost effective and sequence with higher throughput [19]. Third, researchers are beginning to use highly-specific reference genomes that better represent diverse populations and ethnic groups [2, 20, 21, 22, 23, 24, 25, 26]. This may result in the need to map each read set to *multiple* reference genomes that represent various populations within the same species in order to correctly identify the genome donor’s genetic variations (i.e., differences from the most relevant reference genome).

To reduce the large overhead of *fully mapping* a read set to a new reference genome, several existing tools [27, 28, 29, 30, 31, 32, 33, 34, 35, 36] can be used to quickly *remap* the reads (i.e., update a read’s alignment location from the original (old) reference to another (new) reference). In the remainder of this paper, we collectively refer to such methods as *remapping tools*. At a high level, state-of-the-art remapping tools rely on *chain files* (described in Supplementary Section S2), which identify and list *constant regions*, i.e., genome sequences that appear in both old and new references (e.g., regions A and B in Figure 1) and their positional offsets into each reference genome. A remapping tool uses a chain file to identify reads whose original mapping locations in the old reference is sufficiently contained within constant regions and quickly updates the alignment location of each read according to how the location of the constant region containing it changes between the old and new references. For example, Read 2 in Figure 1 can be quickly remapped by shifting its location by 5 base pairs from the old reference to the new reference.^1^

**Figure 1:**
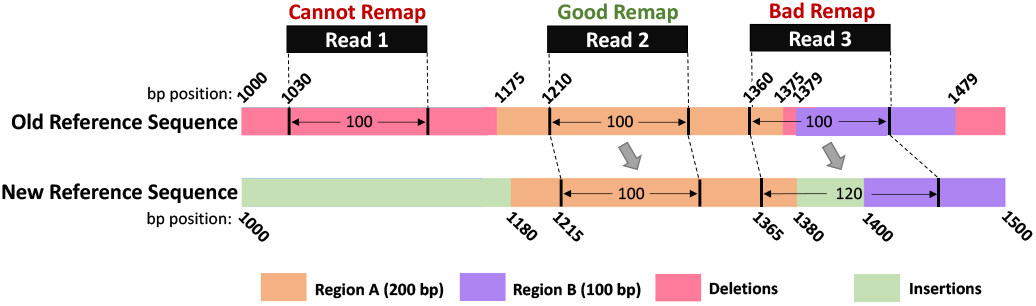
Limitations of Existing Remapping Tools. Existing remapping tools correctly remap reads that mapped *completely within* a region indicated by the chain file (e.g., Read 2). However, these tools 1) cannot remap reads that mapped within a region in the old reference that does not appear in the new reference (e.g., Read 1) and 2) may incorrectly remap reads that align to multiple constant regions in the old reference (e.g., Read 3).

Unfortunately, many these remapping tools 1) are not *comprehensive* in remapping a read set, meaning that they *cannot* remap a significant proportion of reads due to the limitations of using a chain file (e.g., a chain file only contains information about genome sequences that appear exactly the same between two references and their positional offsets into each reference), 2) are not *accurate*, meaning that some remapped reads do *not* align to the sequence they are remapped to in the new reference genome within the acceptable error rate, and 3) result in output on which downstream analysis cannot be performed (i.e., do not provide an end-to-end BAM-to-BAM^2^ remapping solution). We identify two key limitations that we illustrate in Figure 1. First, since each *deleted region* (i.e., a region that does not appear in the new reference) does not have a corresponding region in the new reference, chain files *cannot* provide information on how to remap reads that had originally mapped to a deleted region. This is because, by definition, a deleted region has no similar regions in the new reference. For example, Read 1 in Figure 1 maps to a deleted region in the old reference and therefore cannot be remapped to the new reference to any extent. Second, state-of-the-art remapping tools *only* consider the degree of similarity between a read and the constant regions (from the chain file) in the old reference, without considering the changes in the new reference when remapping the read to the new reference. Therefore, remapping can result in a poor degree of similarity between the read and the new reference. As an example, Read 3 in Figure 1 maps to the old reference with high similarity (i.e., 4 deletions between base pairs 1375 and 1379; < 5% error rate), so it is remapped to the new reference at a location corresponding to the read’s original mapping in the old reference. This remapping does not account for differences that appear in the new reference (e.g., 20 insertions between base pairs 1380 and 1400) and result in a high error rate (i.e., > 5%).

Due to these limitations, existing remapping tools are unable to comprehensively remap a read set from one reference to another. We observe that state-of-the-art remapping tools miss at least 7% of gene annotations when remapping reads from an older human reference genome (hg16) to its latest version (GRCh38), as shown in Supplementary Table S1 and Supplementary Figure S1. These limitations require researchers and practitioners to re-run the *full* genome analysis pipeline for each read set on an updated reference genome for a comprehensive study.

Our **goal** is to provide the first read remapping technique across (reference) genomes 1) that *substantially* reduces the time to remap a read set from an old (i.e., previously mapped to) reference genome to a new reference genome, 2) that is *comprehensive* in remapping a read set, i.e., attempts to remap *all* reads in a read set, 3) provides *accurate* remapping results, i.e., provides alignments with error rates below a specified acceptable error rate, and 4) provides an end-to-end BAM-to-BAM remapping solution on which downstream analysis can be immediately performed. To this end, we propose *AirLift*, the first methodology and tool that leverages the similarity between two reference genomes to satisfy our goal. Specifically, AirLift greatly reduces the time to perform end-to-end BAM-to-BAM remapping on a read set from one reference genome to another while maintaining high accuracy and comprehensiveness that is comparable to *fully mapping* the read set to the new reference. We evaluate AirLift and demonstrate that AirLift satisfies the four design goals of an effective remapping tool by comparing it against state-of-the-art remapping tools and the previous best method of *fully mapping* a read set to a new reference with BWA-MEM [38] across various versions of the human, C. elegans, and yeast references (summarized in Table 1). We demonstrate that AirLift can identify SNPs and Indels with precision and recall similar to full mapping (via GATK HaplotypeCaller [39]) while providing 2.76*×* to 27.4*×* speedup over *fully mapping* a read set to the new reference genome.

**Table 1:** AirLift vs. existing state-of-the-art remapping tools.

| Proposal | Year | Fast | Comprehensive | Accurate | BAM-to-BAM | Memory Usage |
| --- | --- | --- | --- | --- | --- | --- |
| CrossMap [31] | 2014 | ✓ | ✗ | ✗ | ✗ | low |
| LiftOver [27] | 2014 | ✓ | ✗ | ✗ | ✗ | low |
| Full Mapping (BWA-MEM [38]) | 2013 | ✗ | ✓ | ✓ | ✓ | high |
| **AirLift** | 2022 | ✓ | ✓ | ✓ | ✓ | high |

## 2. AirLift

In order to accurately and comprehensively remap a read set, AirLift 1) categorizes and labels each region (i.e., a contiguous sequence within a genome) in the old reference genome depending on its degree of similarity to the most similar region in the new reference and 2) remaps each read from the old reference to the new reference according to the label of the region in the old reference that the read had been originally mapped to.

For each pair of references that AirLift remaps reads between, we must first construct an *AirLift Index*, i.e., a set of lookup tables (LUTs), in a one-time preprocessing step. AirLift queries the AirLift Index with a read and its original mapping location in the old reference (from the BAM file) to efficiently identify the region and the label of the region that the read mapped to in the old reference. This information is then used to identify potential mapping locations of the read in the new reference (based on regions in the new reference that are similar to the region that the read mapped to in the old reference).

We next define these regions, show how to generate the *AirLift Index*, and then explain how to use the *AirLift Index* to quickly remap a read set with high genome coverage.

### 2.1. Reference Genome Regions

We identify four categories of regions that fully describe the relationship between two reference genomes, old and new (shown in Figure 2):

**Figure 2:**
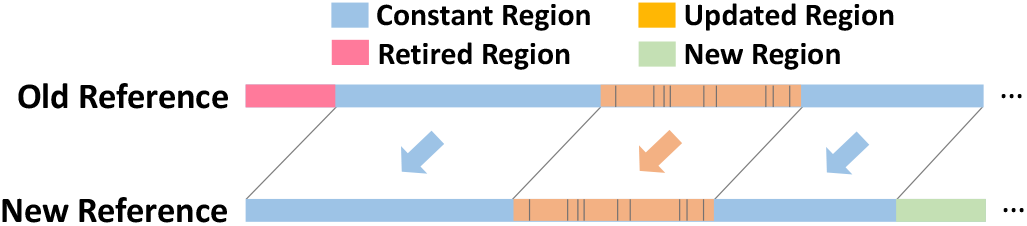
An example pair of reference genomes (old and new) with regions labeled (as constant, updated, retired, and new regions) and associated with each other according to their degrees of similarity. Regions that are associated with (i.e., similar to) each other are indicated with an arrow. Example differences across associated updated regions are shown with black vertical bars.

1. A *constant region* is a region of the genome which is exactly the same in both old and new reference genomes (colored in blue). The start and end positions of a constant region are not necessarily the same in the old and new reference genomes.
2. An *updated region* is a region in the old reference genome that maps to at least one region in the new reference genome within a reasonable error rate, i.e., differences from the old reference (colored in orange with some differences marked with black bars).
3. A *retired region* is a region in the old reference genome that does *not* map to any region in the new reference genome (colored in pink).
4. A *new region* is a region in the new reference genome that does *not* map to any region in the old reference genome (colored in green).

We next describe how we identify and use these regions to quickly and comprehensively remap a read set.

### 2.2. The AirLift Index

The *AirLift Index* is comprised of two lookup tables (LUTs), each of which has a one-time construction cost for any pair of reference genomes. The LUTs describe regions of similarity between a pair of reference genomes, which can then be used to quickly remap reads between the references.

The first LUT, i.e., *constant regions LUT*, associates each constant region in the old reference with its respective region in the new reference genome. AirLift queries this *constant regions LUT* with a location (of a previously-mapped read) in the old reference to quickly find a list of corresponding locations in the new reference that have the same genome sequence. AirLift uses this list of locations to update the mapping of the read, as we explain in more detail in Section 2.4.

The second LUT, i.e., *updated regions LUT*, associates each updated region in the old reference with its respective region in the new reference genome. AirLift queries this *updated regions LUT* with a location (of a previously-mapped read) in the old reference to quickly find a list of corresponding locations in the new reference that have similar genome sequences. AirLift uses this list of locations to update the mappings of the read, as we explain in more detail in Section 2.4.

Once constructed, the *AirLift Index* is used to aid in the efficient mapping of any number of reads from one reference genome to another reference genome. We next explain how to label regions in the reference and construct the *AirLift Index*.

### 2.3. Categorizing Regions of Similarity and Constructing the AirLift Index

The *AirLift Index* is constructed via eight key steps, as we show in Figure 3.

**Figure 3:**
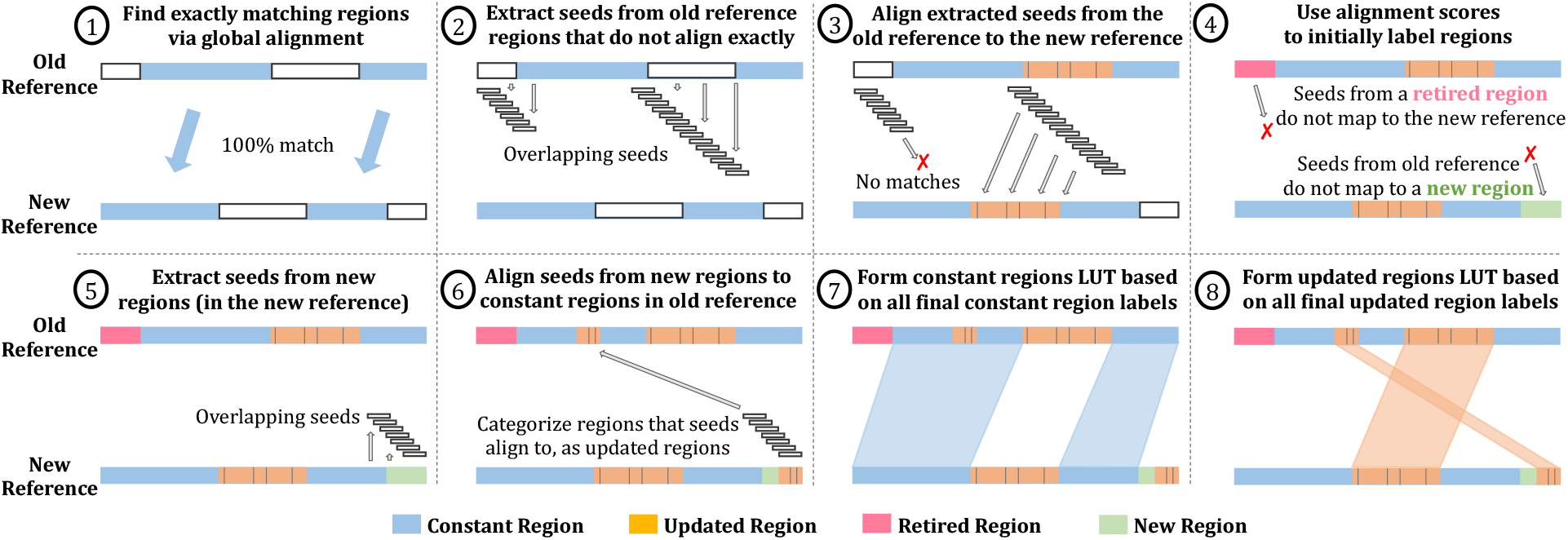
AirLift uses eight key steps to identify and label regions in the old and new reference genomes as *constant, updated, retired*, or *new* in order to efficiently map any number of reads from an old reference genome to a new reference genome.

1. First, we want to identify all regions (i.e., genome sequences) that appear exactly the same in both the old and the new reference genomes. To do so, we use a chain file (described in Supplementary Section S2), which can be generated via BLAT [40] with exact matching (no errors allowed) global alignment. In Figure 3, we indicate the constant regions in blue.
2. In order to label the remaining regions in the new reference, we first extract seeds (i.e., smaller subsequences) from regions in the old reference that do *not* map exactly to the new reference (non-blue regions). Note that these seeds **a)** are the same length (*N*) as the reads that we want to remap, and **b)** are overlapping seeds, i.e., completely overlap with each other such that a seed begins at each base pair within each (non-blue) region and starting *N* – 1 base pairs before each (non-blue) region. This is to ensure that AirLift completely accounts for all possible mapping locations including sequences that may be partially included in a constant region.
3. Next, we map the extracted seeds (from Step 2) to the new reference genome to identify regions of approximate similarity across the reference genomes. Note that this step can be done with *any* read mapper. We label as an *updated region* (colored in orange) 1) any continuous segment of base pairs that any seed has mapped to in the new reference or 2) any continuous segment of seed locations in the old reference whose seeds have mapped to the new reference. Since it is an approximate mapping, we indicate differences between the updated regions in Figure 3 with black stripes. These differences are accounted for by the resulting chain file.
While we describe in more detail how we use these regions in Section 2.4, we can quickly tell that if a read mapped to an updated region in the old reference genome, there is a high chance that the read will map to the respective updated region in the new reference genome. In order to comprehensively identify all possible locations in the new reference that a read can map to just by examining the read’s mapping location in the old reference, we map seeds from the new reference using an error rate of 2*e*, where *e* is the acceptable error rate for a successful alignment, and report the best alignment. Due to our usage of a conservative error rate (2*e*), we are still able to find every potential mapping with an alignment score within the acceptable error rate (Explained in Supplementary Section S4).
4. We find regions in the old reference where seeds (extracted from Step 2) do *not* align to and label them as *retired regions*, since the region or anything similar does not exist in the new reference genome. Similarly, we find regions in the new reference whose seeds do not map to the old reference genome and label them as *new regions*, since the region or anything similar to the region does not exist in the old reference genome.
5. Next, we check to see whether regions within the new regions can be approximately aligned to constant regions in the old reference, since we had *only* previously attempted mapping seeds from non-constant regions to the new regions (in Step 3), and constant regions were only identified with *exact* matching. We do this by first extracting overlapping seeds from the new regions.
6. We then map the extracted overlapping seeds (from Step 5) to the constant regions in the old reference genome. For any seeds that result in a successful alignment, we 1) additionally label the corresponding segment of the constant region as an updated region and 2) relabel the corresponding segment of the new region as an updated region. We can now consider each of these regions as updated regions, since this step has resulted in identifying an associated similar region in the other reference. This step is necessary to ensure that all regions in the old reference are checked for similarity to all regions in the new reference, enabling a comprehensive mapping for reads that map to *any* region in the old reference.
7. We show the associated constant regions between the two references within the areas shaded in blue and use this information to create a *constant regions LUT*, which can be queried with a location in the old reference to obtain locations in the new reference that contain the exact same sequence. We encode the mapping with the chain file format (described in Supplementary Section S2).
8. We show the associated updated regions between the two references within the areas shaded in orange and use this information to create the *updated regions LUT*, which can be queried to immediately return candidate locations in the new reference that a read should be aligned to. We encode the mapping and account for the minor differences using the chain file format.

### 2.4. Using AirLift to Remap a Read

AirLift follows the procedure illustrated in Figure 4 to comprehensively and accurately remap a read set. AirLift first identifies the label of the region that the each read had originally mapped to in the old reference using a series of steps (described in Section 2.4.1). Depending on the label, AirLift remaps each read using one of four independent cases (described in Section 2.4.2), depending on the label of the region that the read originally mapped to within the old reference: **(1)** a read that mapped to a *constant region*, **(2)** a read that mapped to an *updated region*, **(3)** a read that mapped to a *retired region*, and **(4)** a read that *never mapped* to any location in the old reference genome (i.e., an unmapped read).

**Figure 4:**
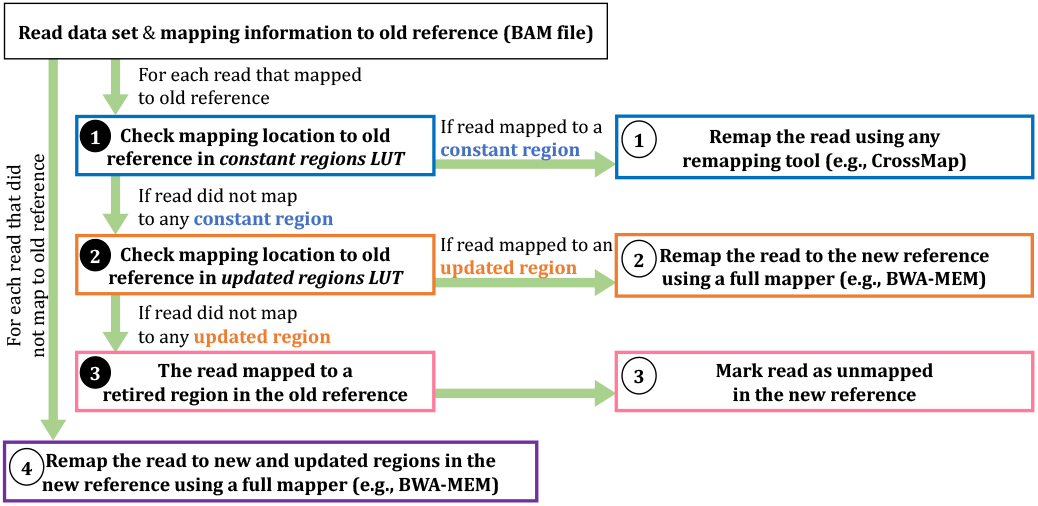
Using AirLift to remap a read set. AirLift remaps each read differently depending on the label of the region in the old reference that the read had originally mapped to: constant, updated, retired, or unmapped.

#### 2.4.1. Determining how to Remap each Read

To determine which case AirLift should apply when remapping a read, AirLift performs the following steps on each read in the read set that originally mapped to any location in the old reference. First, AirLift checks the read’s mapping location to the old reference in the *constant regions LUT* (❶in Figure 4). If the mapping location returns an associated location in the new reference, the read had been originally mapped to a constant region in the old reference and AirLift remaps the read via Case **➀** (described in Section 2.4.2).

If the *constant regions LUT* does not return a location in the new reference, AirLift next checks the read’s mapping location to the old reference in the *updated regions LUT* (❷ in Figure 4). If the mapping location returns an associated location in the new reference, the read had been originally mapped to an updated region in the old reference and AirLift remaps the read via Case **➁** (described in Section 2.4.2).

If the *updated regions LUT* does not return a location in the new reference, the read had been originally mapped to a retired region in the old reference (❸in Figure 4). This is because an old reference is only comprised of constant, updated, and retired regions, and AirLift already determined that the read was not originally mapped to a constant or updated region. AirLift handles such reads via Case **➂** (described in Section 2.4.2).

In order to be comprehensive in remapping a read set, AirLift also considers the reads that were unmapped in the old reference and attempts to remap them to the new reference using Case **➃** (described in Section 2.4.2).

#### 2.4.2. Remapping each Read

**Case 1:** For a read that had originally mapped to a *constant region*, we simply translate the mapping locations according to the offset in the specific constant region from the old reference to the new reference. Since this is the extent of existing state-of-the-art remapping tools capabilities, we can perform this step with any of these tools (e.g., *LiftOver, CrossMap*) for any read that is fully encapsulated within a chain file interval. For our analysis, we built a new tool based on CrossMap that is publicly released called FastRemap [41, 42]^3^, that outputs BAM files which can be used for downstream analysis (e.g., variant calling) for validating our results. The chain file represents only regions that are exact matches, so remapped reads will perfectly match to regions in the new reference genome as well.

**Case 2:** For a read that maps to an *updated region*, we first query the *updated regions LUT* to quickly obtain a list of locations in the new reference genome that are similar (within a 2*e* error rate) to the location that the read mapped to in the old reference genome. We can then use any aligner to align the read to all locations returned by the *updated regions LUT* and return the locations in the new reference genome that align with an error rate smaller than a user defined error rate (*e*).

**Case 3:** For a read that maps to a *retired region* (in the old reference genome), we already know that the read will not map anywhere in the new reference genome, since retired regions are not similar to any region in the new reference genome. Therefore, we can mark that read as an unmapped read in the new reference genome.

**Case 4:** For a read that *never mapped anywhere* in the old reference genome, we know that the read will not map to any constant region in the new reference genome. However, there is a chance that the read can align to updated or new regions in the new reference genome. Therefore, we must fully map the read to each new and updated region using any read mapper.

## 3. Evaluation

Before showing our evaluations of AirLift’s execution time (in Section 3.2), memory usage (in Section 3.3), and accuracy and comprehensiveness (in Section 3.4), we describe our methodology for evaluation.

### 3.1. Evaluation Methodology

#### AirLift Tools

We evaluate AirLift using 1) our new tool based on *CrossMap* [28, 31] that is publicly released and called FastRemap [41, 42], to quickly move all reads that map to constant regions in the old reference and 2) *BWA-MEM* [38] to map reads when constructing the *AirLift Index* and when fully mapping all other reads that do not map to constant regions (i.e., reads that map to updated regions and never mapped to the old reference), according to the *AirLift Index*.

#### Evaluated Remappers

We evaluate two widely used remappers, *CrossMap* [28, 31] and *UCSC LiftOver* [27] to compare against AirLift. Note that these two remappers do *not* provide a *comprehensive* or *accurate* solution to remapping reads from one reference to another. Due to the limitations of prior remappers (described in Supplementary Section S3), we evaluate and compare against the only comprehensive and accurate baseline of *fully mapping* the read set (from scratch without using any prior mapping information) to the new reference genome with *BWA-MEM* [38].

#### Evaluated Reference Genomes

We evaluate AirLift with several versions of reference genomes of varying size across 3 species (i.e., human, C. elegans, yeast) as shown in Supplementary Table S2.

#### Evaluated Read Data Sets

We use DNA-seq read sets from four different samples of the set of species whose reference genomes we examine (as shown in Supplementary Table S3).

#### GATK Variant Calling Evaluation

We evaluate AirLift remapping results via variant calling with *GATK HaplotypeCaller* [39] by following the best practices [43], VCFtools [44] to filter variant calling files based on a minimum quality score of 30 (i.e., ––minQ 30), and use the *hap*.*py* tool (https://github.com/Illumina/hap.py) to benchmark the variant calling results.

#### Evaluation System

We run AirLift on a server with 64 cores (2 threads per core, AMD EPYC 7742 @ 2.25GHz), and 1TB of the memory. We assign 32 threads for C. elegans and yeast and 48 threads for human genomes when running tools with multithreaded capabilities (i.e., SAMtools, BWA-MEM; described in Supplementary Section S7) and collect their runtimes (usr and sys) and memory usage using the `time` command in Linux with `-vp` flags. We report the aggregate runtime (in seconds) and peak memory usage (in megabytes) across all active threads in our evaluations with these configurations.

#### AirLift Evaluation Plots

In each AirLift evaluation plot, we show on the x-axis, both the old reference genome (below) and the new reference genome (above) used in the evaluation. Note that in our evaluations of AirLift, we *only* consider the remapping stage (as other stages are preprocessing stages that are performed once for each pair of reference genomes). We show the execution times of the preprocessing stage in Supplementary Table S4.

### 3.2. AirLift Execution Time

We first demonstrate how AirLift reduces the time to map a set of reads to an updated reference genome by reducing the number of reads that we must map. Figure 5 plots the execution times (y-axis) for mapping a read set to a new reference genome using three different remapping tools, *CrossMap*, AirLift, and *LiftOver* compared to the baseline of *fully remapping* the entire read set from an old reference genome to the new reference genome. We provide the speedup of AirLift over fully mapping the read set to the new reference (i.e., *T*_Full Mapping_/*T*_AirLift_) above each bar.

**Figure 5:**
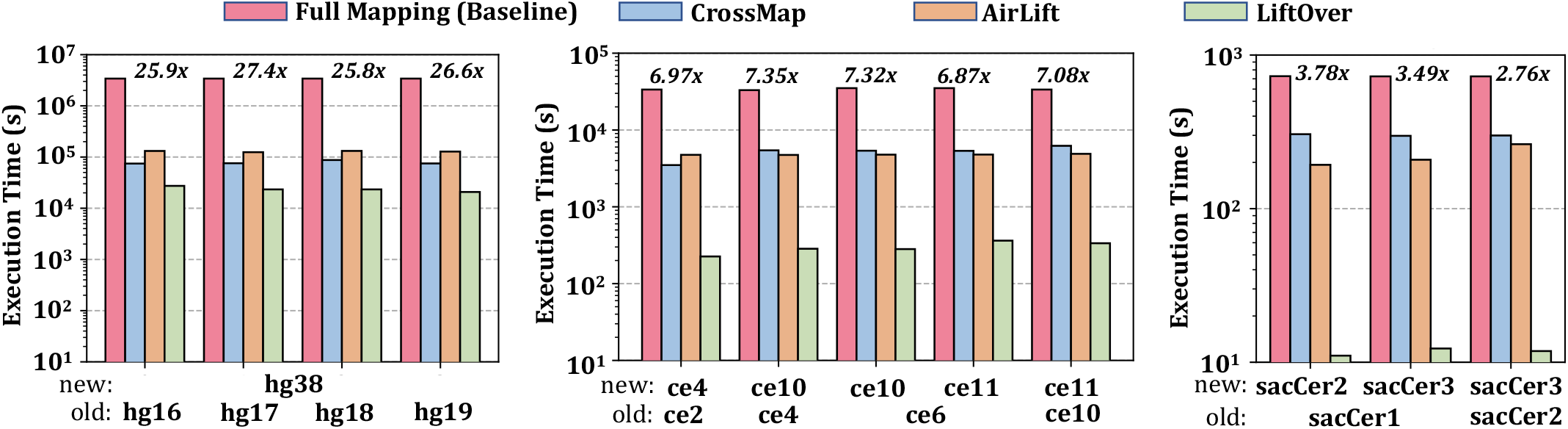
AirLift execution time results. We show the execution time (log-scale y-axis) of running three remapping tools, *CrossMap* (blue), AirLift (orange), and *LiftOver* (green) on a read set to a new reference genome against the baseline (red) of *fully mapping* a read set to the new reference genome. We plot the execution times of each tool for various pairs of reference genomes (x-axis; where the old reference is at the bottom and the new reference is above the old reference) in three separate plots for different sizes of reference genomes, i.e., large (human), medium (C. elegans), small (yeast). We indicate the speedup of AirLift against the *full mapping* baseline above each grouping of bars, since AirLift and the baseline are the only comprehensive and accurate remapping techniques available.

The execution time of AirLift is calculated as the sum of the execution times for performing each of the cases (described in Section 2.4.2) as shown in Equation 1, where *T*_constant_ is the time to translate all reads that originally map to a constant region in the old reference, *T*_updated_ is the time to map all reads that originally mapped to an updated region in the old reference, *T*_retired_ is the time to map all reads that originally mapped to a retired region in the old reference, and *T*_unmapped_ is the time to map all reads that never mapped anywhere in the old reference. The exact execution time breakdowns for each of these four cases are shown in Supplementary Table S6). We also provide the number of reads that AirLift must remap in each case for each pair of references in Supplementary Table S7, and the average time per read per case for each pair of references in Supplementary Table S7.

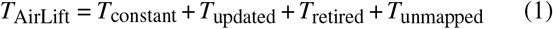

We make three observations based on Figure 5 and the supplementary tables. First, AirLift consistently provides significant speedup over the baseline (of *fully mapping* a read set) across all tested pairs of references, ranging from 2.76 *×* (`sacCer2`→`sacCer3`) up to 27.4 *×* (`hg17 hg38`). Second, AirLift execution time is largely comprised of the time to remap reads that originally mapped to the constant region in the old reference. This is because the number of reads remapped by AirLift are mostly (i.e., between 86.57% for `hg16`→`hg38` and 98.47% for `ce10`→`ce11`) comprised of reads that originally mapped to a constant region. Third, AirLift execution time is significantly lower than the full mapping baseline since the average time to remap a read from the constant regions is significantly lower than the average time to fully map a read. This is because AirLift can very efficiently remap reads from constant regions. Fourth, remapping a read set with AirLift between a pair of references with a smaller constant regions size results in a higher execution time. Therefore, AirLift performs faster when remapping reads between pairs of references that are more similar to each other.

We conclude that AirLift significantly improves the execution time for *comprehensively* and *accurately* remapping a read set from an old reference to a new reference compared to the baseline of *fully mapping* the read set to the new reference.

### 3.3. AirLift Memory Usage

Figure 6 plots the peak memory usage in MB (y-axis) across the remapping tools (i.e., *CrossMap*, AirLift, and *LiftOver*) and baseline full mapping method (i.e., *BWA-MEM*) for our set of evaluated reference pairs (x-axis). We find that across all tested reference pairs, AirLift has similar peak memory requirements as our *full mapping* baseline, *BWA-MEM*. This is because AirLift relies on *BWA-MEM* to remap a portion (i.e., up to 16.61%) of the read set, which is large enough to require the same amount of memory as mapping the full read set.

**Figure 6:**
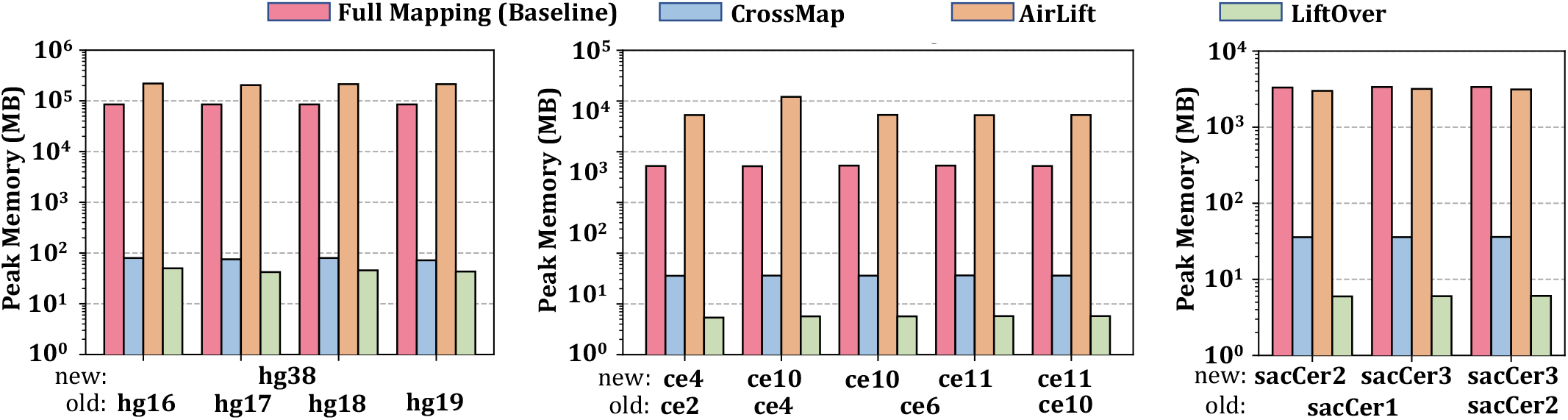
AirLift memory usage results. Peak memory usage results for each of the remapping tools during remapping.

### 3.4. GATK Variant Calling Results

To demonstrate that AirLift provides similar mapping results as a *full mapper* (baseline) and it is much more comprehensive and accurate than *CrossMap* and *LiftOver*^4^, we perform downstream analysis (i.e., variant calling). We use the GATK HaplotypeCaller tool to call variants from both the 1) *full mapping* BAM file and 2) Airlift-generated BAM file. We use the VQSR [45] tool to recalibrate the variants based on quality scores provided by the GATK HaplotypeCaller tool. We use the hap.py tool to benchmark 1) the AirLift variant calls against full mapping, 2) the AirLift variant calls against the gold standard (i.e., ground truth), and 3) full mapping variant calls against the ground truth, if the ground truth is available. We use the variant calling ground truth from the Platinum Genomes [46] and Genome in a Bottle [47] for the human NA12878 sample. We only benchmark Airlift against *full mapping* for the C. elegans and yeast data sets, since we do not have the ground truth for these species. We report the precision and recall results for the SNPs and insertion-deletions (indels) as calculated by hap.py (https://github.com/Illumina/hap.py).

Table 2 shows the variant calling results for human, C. elegans, and yeast genomes, respectively. Each row contains quality measurements of identifying single nucleotide polymorphisms (SNPs) and insertion-deletions (indels) for a pair of reference genomes in terms of precision and recall (written as ‘precision score(%)/recall score(%)’). For the human results, we show the precision and recall scores of *full mapping* when identifying the set of SNPs and indels compared against the set of SNPs and indels that the ground truth reports, to demonstrate how AirLift compares against *full mapping* when identifying ground truth SNPs and indels. The columns are separated to show separate precision and recall scores for identifying the set of SNPs and indels when compared against the set of SNPs and indels that *full mapping* identifies (vs. Full Mapping) and the ground truth reports (vs. Ground Truth; only available for human results).

**Table 2:**
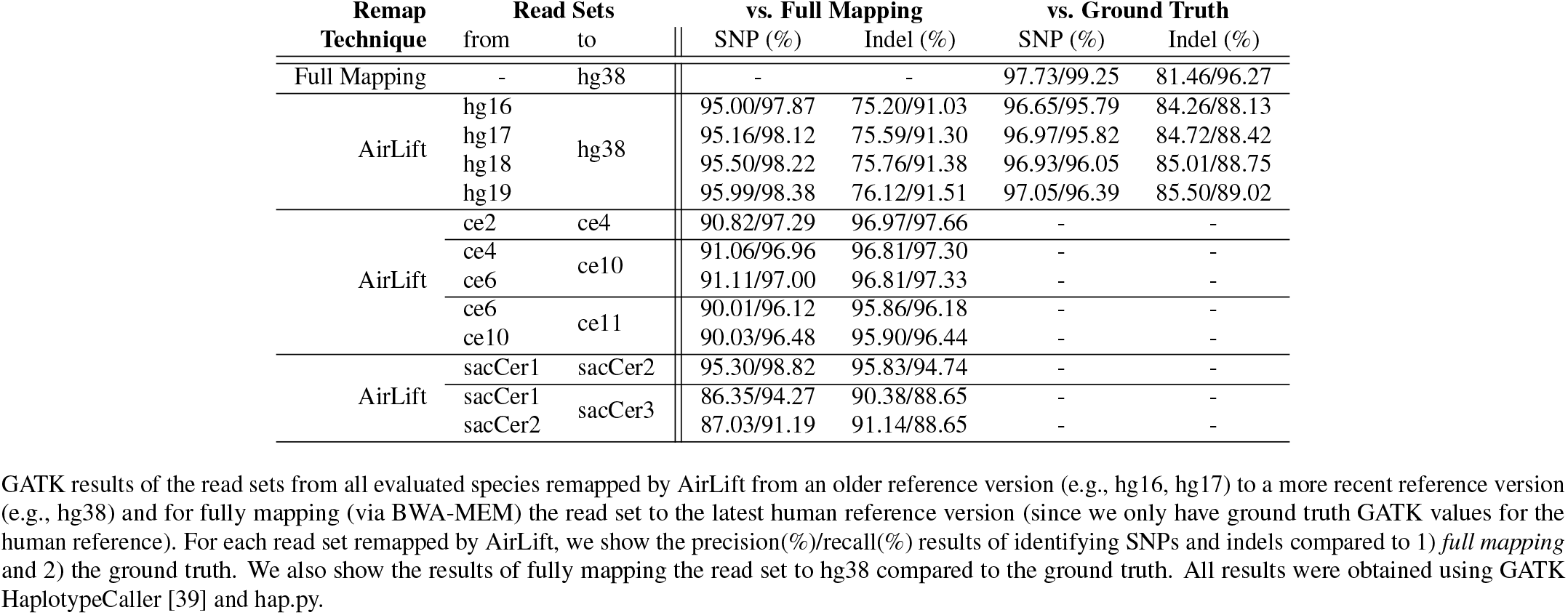
GATK Variant Calling Results for Human, C. elegans, and Yeast Genomes.

We make two key observations. First, we observe that AirLift is able to identify SNPs reported by *full mapping* with high precision and recall scores (as shown under the first column, *vs. Full Mapping*). This is because AirLift 1) identifies all possible mapping locations for each read in the read set similarly to the *full mapping* approach, 2) comprehensively maps each read accordingly, and 3) reports accurate alignment results (i.e., alignments with error rates below a specified acceptable error rate) unlike existing remapping tools. Second, we observe that AirLift indentifies SNPs and indels reported by the ground truth with precision similar to *full mapping*. We observe this by comparing the results in the first row (i.e., *Full Mapping*) against the AirLift results directly underneath them (e.g., 97.73%/99.25% precision/recall values for identifying SNPs when full mapping to hg38 compared to 97.05%/96.39% when using AirLift between `hg19`→`hg38`; only available for human results). We note the small variation across precision and recall values in the table and attribute them to two main factors. The first factor is due to the recalculation of mapping quality scores when fully mapping reads whereas AirLift may report mapping results that do not necessarily result in the best alignment score, since AirLift performs the most efficient method for remapping a read in the case that multiple regions are available (i.e., a read that maps to a constant region and updated region will be treated as a read in a constant region). Fully recalculating mapping scores enables improving the overall accuracy of the mapping quality in an alignment, which improves the overall accuracy in variant calling [48]. This is a trade-off between using an efficient and highly accurate remapping tool, AirLift, and fully mapping reads with slightly higher accuracy that comes with significantly higher computational costs. The second factor is due to the discrepancies that may occur as a result of genomic repeats and reproducibility issues in BWA-MEM [49]. We argue that these alignment differences do not cause a significant loss in variant calling quality, as AirLift precision and recall results for SNPs and indels are very similar to full mapping (when both are benchmarked against the ground truth).

We have shown in our evaluations against existing stateof-the-art remapping tools, that AirLift can comprehensively and accurately remap a read set from one reference genome to another at high speeds (i.e., up to 27.4*×* faster than our *full mapping* baseline). Since AirLift accomplishes our four goals of remapping a read set quickly, comprehensively, accurately, and end-to-end, providing a BAM-to-BAM result that can be immediately used in downstream analysis, we conclude that AirLift is a viable tool to be used as a quick alternative to fully mapping a read set when it had previously been mapped to a similar reference genome.

## 4. Discussion

We briefly discuss AirLift’s limitations and several directions for future work.

### Limitations

We identify two key limitations of AirLift. First, the AirLift index must be produced with several parameters. This limitation requires users to rebuild an AirLift Index each time the remapping parameters (e.g., read size, alignment heuristics) change. Due to the high one-time cost of generating an AirLift Index, users that remap large read data sets (with the same parameters) between two reference genomes are not affected. However, it may limit the performance benefit when remapping many read sets with varying parameter requirements or to many different reference genomes. Second, AirLift provides high accuracy in identifying SNP/INDEL variants. Although it is lower than fully mapping a read data set, we believe that AirLift provides very high speedup relative to its cost in accuracy when comparing against full mapping.

### Future Work

Addressing the key limitation of AirLift to any degree would significantly expand its applicability to various use cases. However, due to the complexity of implementing such algorithmic features, we leave this to future work. We believe that despite its current limitations, AirLift advances the state-of-the-art and provides a good first step into the design of fast and comprehensive remapping tools.

## 5. Conclusion

We introduce AirLift, a methodology and tool for quickly, comprehensively, and accurately remapping a read data set that had previously been mapped to an older reference genome to a newer reference genome. AirLift is the first tool that provides BAM-to-BAM remapping results of a read data set on which downstream analysis can be immediately performed. The key idea of AirLift is to construct and use an *AirLift Index*, which exploits the similarity between two references to quickly identify candidate locations that a read should be remapped to based on its original mapping in the old reference. We compare AirLift against several existing remapping tools, CrossMap and LiftOver, which we demonstrate have several major limitations. These tools either do *not* provide accurate and comprehensive remapping results or do not result in remapping results on which downstream analysis can be immediately performed (summarized in Table 1). We compare AirLift against the only comprehensive and accurate method of *fully mapping* a read data set to the new reference using BWA-MEM, and find that AirLift significantly reduces the execution time by up to 27.4 *×*, 7.35 *×*, and 3.78*×* for large (human), medium (C. elegans), and small (yeast) reference genomes, respectively. We validate our results against the ground truth and show that AirLift identifies similar rates of SNPs and Indels as the full mapping baseline. We conclude that AirLift is the first comprehensive and accurate remapping tool that substantially reduces the execution time of remapping a read data set, while providing end-to-end BAM-to-BAM results on which downstream analysis can be performed. We look forward to future works that take advantage of as well as improve AirLift for various genomic analysis studies.

## Supporting information

Supplemental Pages

## Data Availability

The Human NA12878 illumina read data set is publicly available (Accession number ERR194147 and ERR262997). The C. elegans N2 illumina read data set is publicly available (Accession number SRR3536210). The Yeast S288C illumina read data set is publicly available (Accession number ERR 1938683).

## Supplementary Text for

### S1. Currently Available Remapping Tools

#### UCSC LiftOver

One of the most commonly used remapping tools is UCSC LiftOver [1]. UCSC LiftOver uses a chain file between two different assemblies of a genome to convert the coordinates from one assembly to the assembly of the other genome. UCSC LiftOver suffers from three major shortcomings. First, UCSC LiftOver functionality is limited to the genomes whose assemblies are provided by the UCSC Genome Browser [2], hence, making it impossible to remap genomes whose assemblies are not yet included in the tool. Second, the tool *only* converts the coordinates of regions within the old reference genome that are highly similar to regions within the updated reference genome and ignores regions with significant variance, which prevents a comprehensive remapping of the coordinates (described in more detail in Supplementary Section S3). Third, UCSC LiftOver only supports BED-format (i.e., browser extensible data) input files which limits its usage even further.

#### CrossMap

One alternative to UCSC LiftOver is CrossMap [3, 4]. CrossMap follows a similar approach with UCSC LiftOver and uses chain files to convert mappings from an older reference genome to a newer reference genome. Compared to UCSC LiftOver, CrossMap supports a larger set of input file formats, such as BAM, SAM, or CRAM, BED, Wiggle, BigWig, GFF (i.e., general feature format) or GTF (i.e., gene transfer format), and VCF (i.e., variant call format) [3, 4]. Unfortunately, CrossMap suffers from similar limitations as UCSC LiftOver.

#### NCBI Genome Remapping Service

Another alternative is NCBI Genome Remapping Service [5], which also remaps the annotations from one genome assembly to another. NCBI Remap has support for a larger set of input/output file formats, such as BED, GFF, GTF, and VCF. NCBI Remap can also perform cross species remapping for a limited number of organisms. However, as with UCSC LiftOver, NCBI Remap is limited by the provided assemblies.

#### Segment_liftover

Segment_liftover [6, 7] is another tool that is designed to map coordinates of one genome assembly to another genome’s assembly while maintaining the integrity of the genome segments that are not continuous anymore in the target assembly. However, Segment_liftover first runs UCSC LiftOver and then attempts to approximately map any failed conversions [6]. Due to the high coverage of UCSC LiftOver in remapping segments, most conversions are performed by UCSC LiftOver and therefore suffer from the same shortcomings of UCSC LiftOver.

#### Galaxy

Galaxy [8, 9] is a web-based platform, which has LiftOver as part of its toolset. This tool is based on UCSC LiftOver [1] and the chain files provided by UCSC Genome Browser [2]. Thus, Galaxy also suffers from similar limitations as UCSC LiftOver.

#### PyLiftover

PyLiftover [10] is a Python implementation of a limited version of UCSC LiftOver. PyLiftover does not convert ranges (i.e., only converts point coordinates) between different assemblies, and it does not support BED-format input files.

#### Bazam

Bazam [11] is another tool which remaps short paired reads by optimizing memory usage while providing high parallelism. However, Bazam *only* targets the steps where reads are read from a BAM or CRAM file (i.e., read extraction) and sent to an aligner (e.g., BWA [12]). Eventually, *all* the reads are remapped to the new reference genome, which is inefficient.

#### nf-LO

nf-LO [13] is a containerized implementation of UCSC LiftOver written in Nextflow, which enables its usage in any Unix-based system. However, as nf-LO is directly based on UCSC LiftOver, it comes with the same limitations.

#### LevioSAM

LevioSAM [14, 15] is a remapping tool that remaps reads from a variant-aware reference to another reference using a VCF file or a chain file. LevioSAM creates a separate index file for querying remapping and updating the CIGAR string of remapped reads. AirLift provides an efficient and comprehensive lookup for reads that can be remapped without updating the CIGAR string and sends the remaining reads to a read mapper to perform a full read mapping, which can align these reads to better regions with updated CIGAR strings.

#### Liftoff

Liftoff [16] uses similar methods as remapping tools, but focuses primarily on remapping genes between two references. AirLift on the other hand remaps the full read set between two references, providing coverage on non-coding regions of the genome that may contain important information (e.g., gene expression).

### S2. Chain File Format

The chain file [17] is a commonly used data structure across remapping tools and essentially describes the relationship of two reference genomes. The chain file is typically generated with two steps: 1) performing global alignment to detect similar regions between two reference genomes, and 2) encoding the identified similarities into a simple readable format. The chain file encodes the differences of large genomic sequences (e.g., chromosomes) as a list of three-integer-tuples. The first integer represents the length of the *alignment strand*, or shared sequence. The second integer represents the length of the *gap*, or different sequence in the old reference genome. The third integer represents the length of the gap in the new reference genome. In this way, the offset of an alignment strand across the old and new reference genomes can be quickly calculated, and reads that fall within the alignment strand can be quickly remapped according to the offset.

### S3. Limitations of Currently Available Remapping Tools

Repeating a genomic study using a different version of the reference genome is computationally very expensive. A faster and more convenient way to achieve this is to “remap” the mapping locations from the older reference genome to its updated version [1, 3, 4, 5, 6, 7, 9, 10]. While these tools (described in Section S1) can quickly move many annotations, there are several limitations with the current methodology that we study and demonstrate with UCSC LiftOver [1]. UCSC LiftOver is both the state-of-the-art tool commonly used for remapping reads and also the codebase wrapped or modeled by several other tools [7, 8, 9, 10]. In Figure 1, read 2 (mapped to location 1210 in the old reference) shows an example of how a read is remapped using such a tool. The tool identifies a region corresponding to the region that read 2 maps to (region A) that is similar across the two reference. Since the region begins at location 1180 in the new reference and to 1175 in the old reference, all reads mapping to region B are also shifted by 5 base pairs when remapping them to the new reference.

In our evaluation of UCSC LiftOver [1], we find that techniques relying on the standard chain file format do not account for large insertions (i.e., many new base pairs that exist in the new reference but not in the old reference) in the new reference genome. Discounting insertions results in two problems when using these techniques: 1) a remapped read can contain a large insertion in the new reference resulting in a poor alignment and low accuracy, and 2) insertions have low coverage in the new reference due to the limitations of chain files resulting in low coverage of the new reference. We illustrate these issues in Figure 1 (Reads 3 and 1, respectively).

#### S3.1. Limitation 1: Low Accuracy

The first limitation we identify is that UCSC LiftOver [1] only accounts for the overlap between a read and alignment sequences in the old reference genome when remapping the read. A read will be remapped to the new reference genome if the total length of gaps in the old reference genome between the start and end of the read is less than the read length multiplied by the selected error acceptance rate (5% is typically used in read alignment. This corresponds to the *Minimum ratio of bases that must remap* parameter on the UCSC LiftOver webtool [1] being set to 0.95). However, the tool remaps the read *regardless* of the total length of the gaps in the new reference genome. This means, that if there is a large insertion in the new reference genome between the start and end of the read in the alignment strand, the read will *still* be mapped even if the read no longer aligns to that location with an error acceptance rate of 5%. For example, read 3 in Figure 1 maps to the old reference genome at location 1360. While there are 4 base pairs of difference (1375-1379) at the read’s mapping location in the old reference, it is within the 5% error acceptance rate and therefore will be remapped to location 1365 in the new reference. The new mapping in the new reference has an insertion of 20 base pairs long, which means that the new mapping can have an error rate of 20% (which is well beyond the 5% error acceptance rate). In our evaluation of UCSC LiftOver (with an error acceptance rate of 5%, and reads of length 100 base pairs), 0.41% of remapped reads resulted in an error rate greater than 5% (often times much greater, i.e., >40%) when aligned to the sequence at the remapped location in the new reference genome.

#### S3.2. Limitation 2: Low Coverage

The second limitation we identify is that UCSC LiftOver [1] is inherently unable to remap reads 1) to regions in the new reference genome with large insertions (i.e., regions that do not appear in the old reference) or 2) that map to deleted regions in the old reference (i.e., regions that do not appear in the new reference). This results in low coverage of those regions in the new reference. For example, Read 1 in Figure 1 maps to the old reference genome in a deleted region (1030-1130). However, since the chain file cannot relay how that region relates to the new reference, the read cannot be moved to the new reference. In addition, the large insertion in the new reference (1000-1180) does not get mapped to since reads never mapped to regions similar to it in the old reference genome. To demonstrate the implications of this limitation, we examine chain files to identify the *theoretical* minimum number of annotations that are missed due to the limitation (i.e., any annotation that falls within regions in the new reference that are not covered by the chain file). In Supplementary Figure S1, we show the minimum amount of information lost when remapping from one human reference genome version (x-axis) to the latest human reference genome version (hg38). The y-axis shows the minimum percentage of annotations (labeled and marked with unique colors) missed when remapping solely with existing chain files. We make two key observations based on Supplementary Figure S1. First, we observe that a significant portion (>7%) of the genes and transcripts are lost when simply using an available remapping tool (i.e., UCSC LiftOver) between hg16 and hg38. Second, the percentage of the missed annotations decreases as the difference in versions becomes smaller, but even when lifting annotations between hg19 (released in 2009) and hg38 (released in 2013), 4.47% of genes are “lost” in hg38. Supplementary Table S1 contains the exact values of each lost annotation (in percentages) when using UCSC LiftOver from hg16, hg17, hg18, and hg19 (rows) to hg19 and hg38 (columns). We expect to observe similar behavior in tools that wrap UCSC LiftOver (e.g., [7, 8, 9, 10]).

#### S3.3. The Need for a Comprehensive Remapping Tool

As the output of lifting annotations from one reference to another is used in downstream genome analysis, we argue that the speed and accuracy of lifting annotations, and coverage of the new reference genome are *all* crucial. However, prior works mainly focus on the speed at the cost of both accuracy and coverage. These remapping tools are often very inaccurate and can only lift mappings or annotations for regions with minor changes [18]. Therefore, if researchers want a comprehensive study using a new reference genome, they must map the entire read data set to the new reference genome rather than rely on the results of such remapping tools [18]. Due to the high similarity between the old and new reference genomes, we can use information from the old mapping to *very quickly* map a read data set to an updated reference genome. **Our goal** is to produce a method for quickly remapping the reads of a sample from one reference genome to an updated version of the reference genome or another similar reference genome with high genome coverage.

### S4. Creating the AirLift Index with a 2*e* Error Rate

In Supplementary Figure S2, we illustrate an example of why using a 2*e* error rate enables AirLift to find all possible alignments of a read in the old reference. In Supplementary Figure S2, a read (of length 20) aligns to a subsequence in the updated region of the old reference genome with an *e* = 5% error rate (one mismatch on the 9^*th*^ base pair), and also aligns to a subsequence in the updated region of the new reference genome with an *e* = 5% error rate (one mismatch on the 16^*th*^ base pair). While the read could map to either of the regions with a 5% (e) error rate, the sequences between the updated regions exhibit a 10% (2e) error rate, and thus we could only identify the new region as a potential match if we use a 2*e* error rate when categorizing regions.

### S5. Prior Tools Compatability with GATK

#### UCSC LiftOver

UCSC LiftOver only generates Browser Extensible Data (BED) files when remapping read sets. BED files are incompatible with variant calling tools, as they lose a lot of information required for variant callers.

#### CrossMap

CrossMap incorrectly handles supplemental alignments, resulting in duplicate mappings in the BAM file that cannot be used for variant calling. We have made a few updates to CrossMap as can be found in the forked repository https://github.com/canfirtina/CrossMap.

### S6. AirLift Index Study

We first analyze the AirLift Indices to determine the breakdown of regions across both old and new references. Supplementary Table S5 shows the region sizes (i.e., constant, updated, retired) that an old reference is comprised of when preprocessed with another reference. We note that the closer the version numbers between the pair of references are to each other, 1) the larger the constant region is, and 2) the smaller the updated region is. This is intuitive as each reference genome version releases incremental changes to update missing and inaccurate sequences, so the similarity between consecutive releases would likely be higher than that between releases further apart. We also observe, as expected, that the percentage of reads that map to a region in the reference is correlated with the region size (i.e., larger regions have more reads mapped to that region).

Since the most expensive method for remapping in AirLift (i.e., full mapping via BWA-MEM) is employed *only* for reads that mapped to updated regions of the old reference or never mapped at all, we can expect a significant reduction in the mapping time, based on the small updated regions of Supplementary Table S5.

### S7. Running AirLift in Multithreaded Mode

If a user specifies multiple threads to execute AirLift, all multithreaded-enabled tools (i.e., SAMtools, BWA-MEM, and FastRemap, our in-house C++ implementation of CrossMap) used in the execution pipeline are executed with the specified number of threads. First, AirLift extracts alignments to the constant regions of the old reference in parallel using *SAMtools view* with the multithreading option (i.e., “–threads”) enabled to copy these alignments to a temporary BAM file called *fastremap_before*.*bam* file. Second, AirLift uses *SAMtools index* to generate the index file for *fastremap_before*.*bam* in parallel (using the “-@” option). Third, AirLift uses FastRemap to update the alignment positions in the *fastremap_before*.*bam* file according to the new reference genome. FastRemap uses the Seqan library, which utilizes as many CPU cores available when executing its function calls. As FastRemap generates the alignments with the updated positions, the alignments are piped for sorting using *SAMtools sort*, which runs in parallel (using the “–threads” option). Fourth, AirLift aligns reads that fall either in the retired or updated regions of the old reference with a read mapper (e.g., BWA-MEM) in multi-threaded mode. Fifth, AirLift uses *SAMtools merge* to combine all intermediate alignment results (i.e., mapped and remapped alignments) in parallel (using the “–threads” option).

## Supplementary Tables for

**Table S1:**
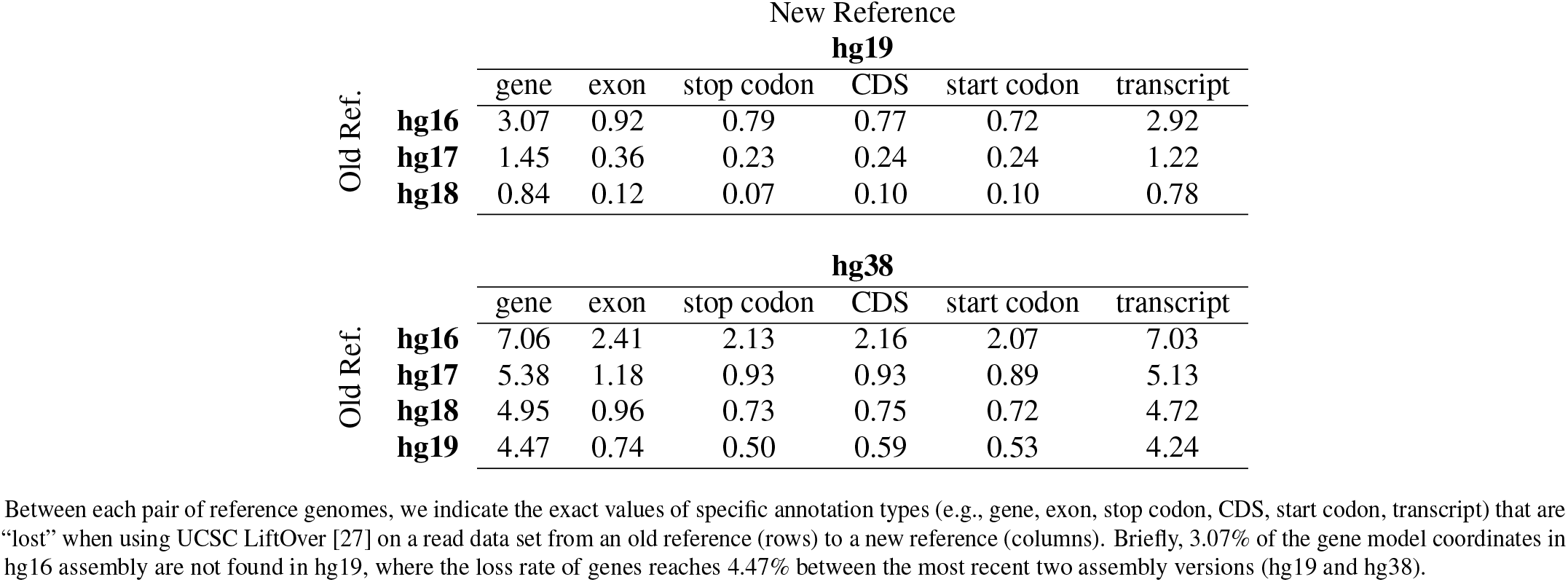
Annotations in the new reference not covered by reads when remapping reads across reference genomes with a remapping tool that solely relies on chain files (e.g., UCSC LiftOver).

| Old Ref. |  | New Reference |  |  |  |  |  |
| --- | --- | --- | --- | --- | --- | --- | --- |
|  |  | **hg19** |  |  |  |  |  |
|  |  | gene | exon | stop codon | CDS | start codon | transcript |
| Old Ref. | **hg16** | 3.07 | 0.92 | 0.79 | 0.77 | 0.72 | 2.92 |
|  | **hg17** | 1.45 | 0.36 | 0.23 | 0.24 | 0.24 | 1.22 |
|  | **hg18** | 0.84 | 0.12 | 0.07 | 0.10 | 0.10 | 0.78 |
|  |  | **hg38** |  |  |  |  |  |
|  |  | gene | exon | stop codon | CDS | start codon | transcript |
| Old Ref. | **hg16** | 7.06 | 2.41 | 2.13 | 2.16 | 2.07 | 7.03 |
|  | **hg17** | 5.38 | 1.18 | 0.93 | 0.93 | 0.89 | 5.13 |
|  | **hg18** | 4.95 | 0.96 | 0.73 | 0.75 | 0.72 | 4.72 |
|  | **hg19** | 4.47 | 0.74 | 0.50 | 0.59 | 0.53 | 4.24 |

Between each pair of reference genomes, we indicate the exact values of specific annotation types (e.g., gene, exon, stop codon, CDS, start codon, transcript) that are “lost” when using UCSC LiftOver [27] on a read data set from an old reference (rows) to a new reference (columns). Briefly, 3.07% of the gene model coordinates in hg16 assembly are not found in hg19, where the loss rate of genes reaches 4.47% between the most recent two assembly versions (hg19 and hg38).

**Table S2:**
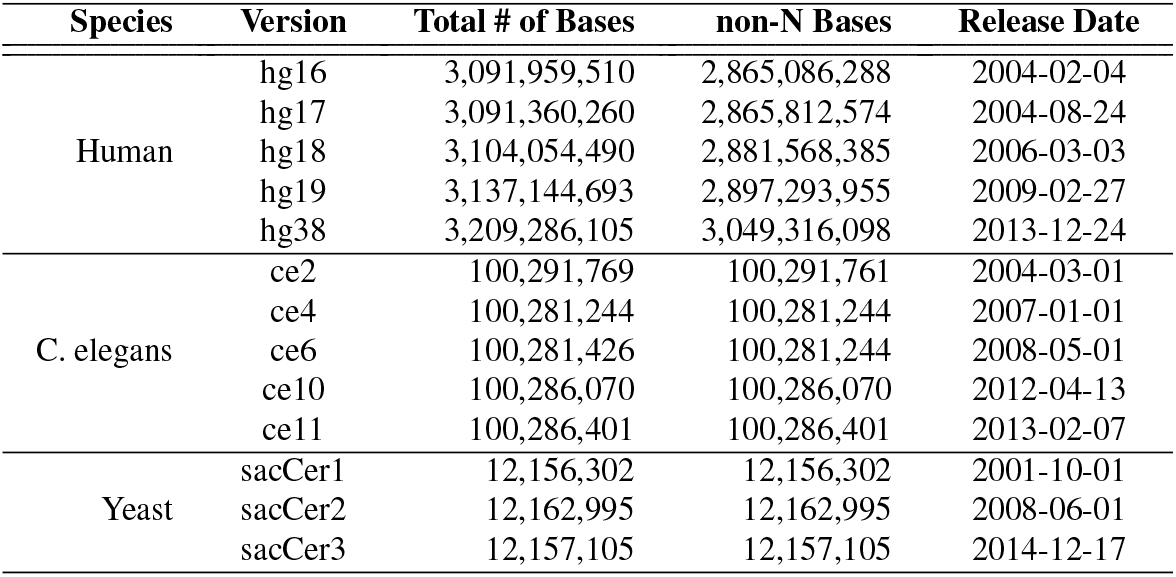
Details of the reference genomes that we use in our evaluations.

**Table S3:**
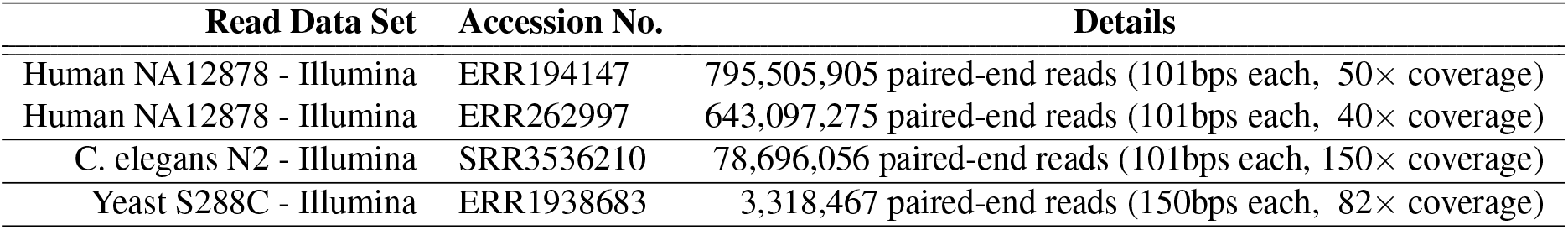
Read data sets that we use in our evaluations.

**Table S4:**
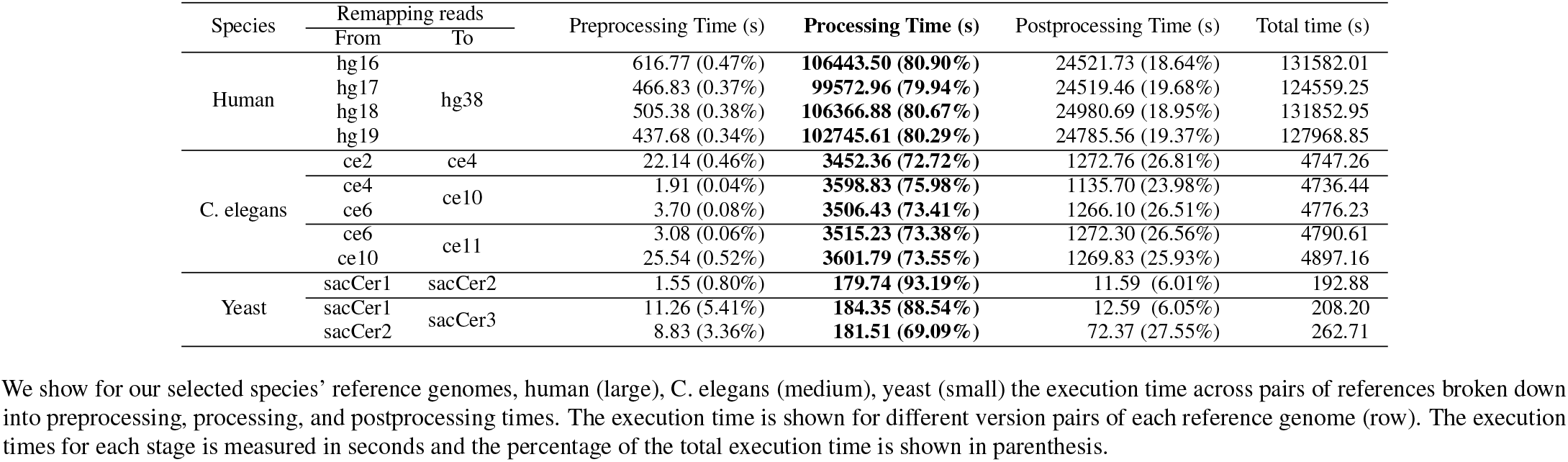
AirLift Execution Time Breakdown.

**Table S5:**
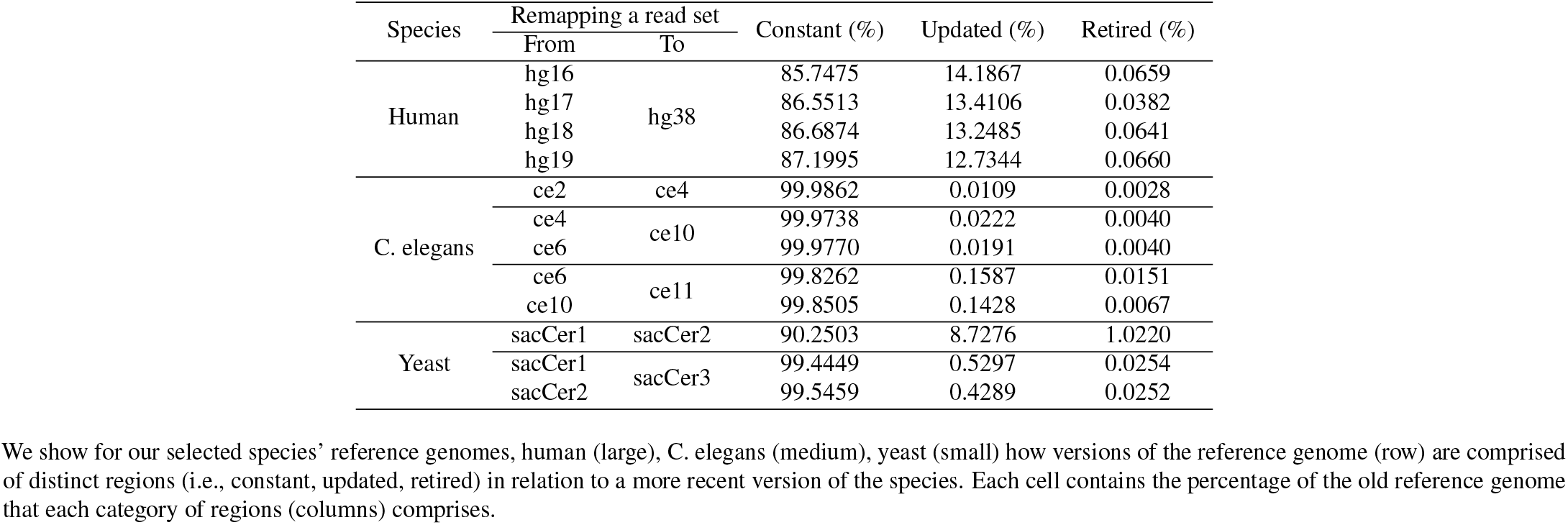
Breakdown of Region Labels for Each Pair of Reference Genomes.

**Table S6:**
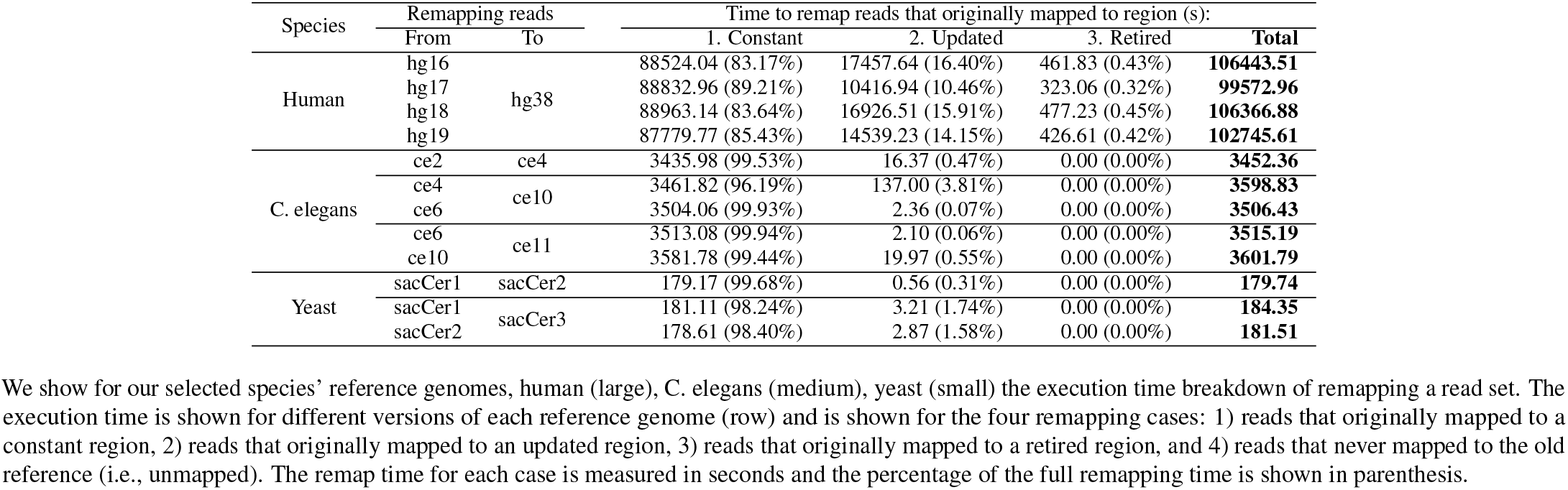
Execution Time Breakdown for Remapping a Read Set by Case.

**Table S7:**
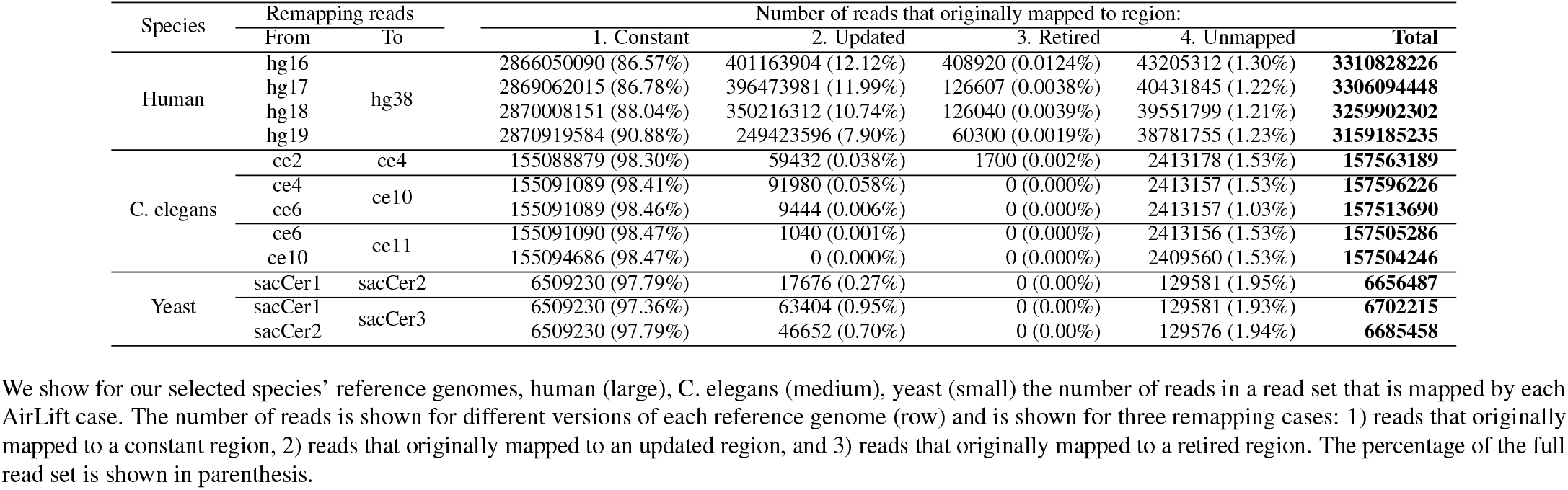
Number of Reads Remapped by Each AirLift Case.

**Table S8:**
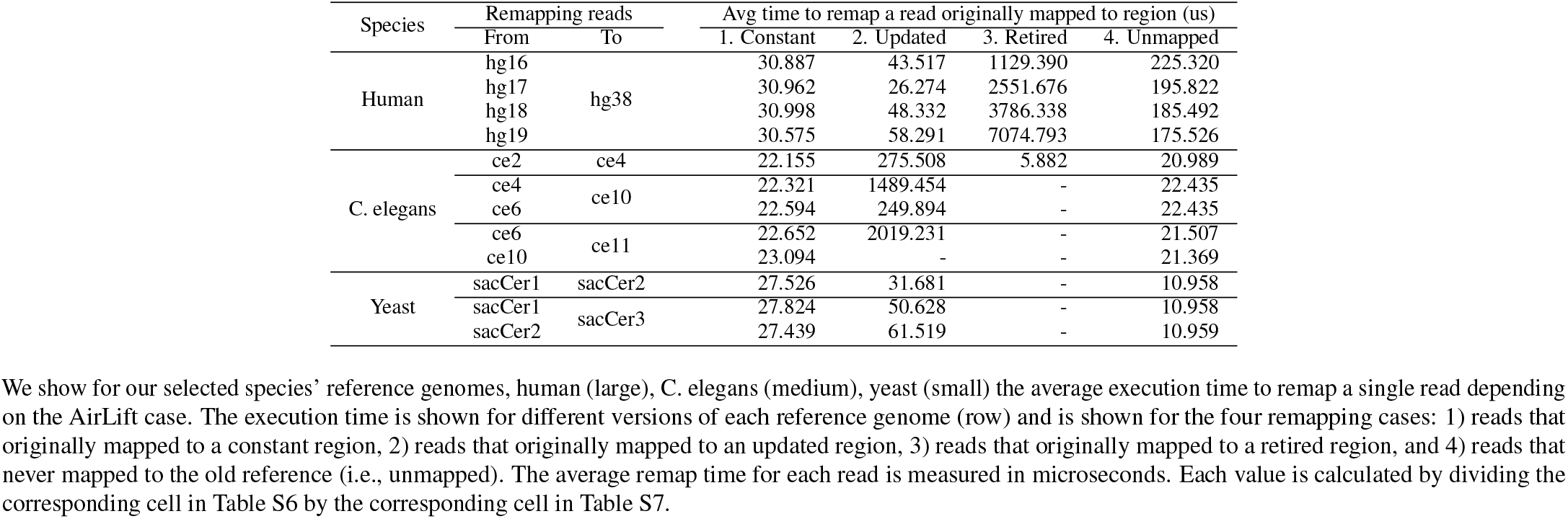
Execution Time per Read when Remapping a Read Set by Case.

| Species | Remapping reads |  | Avg time to remap a read originally mapped to region (us) |  |  |  |
| --- | --- | --- | --- | --- | --- | --- |
|  | From | To | 1. Constant | 2. Updated | 3. Retired | 4. Unmapped |
| Human | hg16 | hg38 | 30.887 | 43.517 | 1129.390 | 225.320 |
|  | hg17 |  | 30.962 | 26.274 | 2551.676 | 195.822 |
|  | hg18 |  | 30.998 | 48.332 | 3786.338 | 185.492 |
|  | hg19 |  | 30.575 | 58.291 | 7074.793 | 175.526 |
| C. elegans | ce2 | ce4 | 22.155 | 275.508 | 5.882 | 20.989 |
|  | ce4 | ce10 | 22.321 | 1489.454 | - | 22.435 |
|  | ce6 |  | 22.594 | 249.894 | - | 22.435 |
|  | ce6 | ce11 | 22.652 | 2019.231 | - | 21.507 |
|  | ce10 |  | 23.094 | - | - | 21.369 |
| Yeast | sacCer1 | sacCer2 | 27.526 | 31.681 | - | 10.958 |
|  | sacCer1 | sacCer3 | 27.824 | 50.628 | - | 10.958 |
|  | sacCer2 |  | 27.439 | 61.519 | - | 10.959 |

## Supplementary Figures for

**Figure S1:**
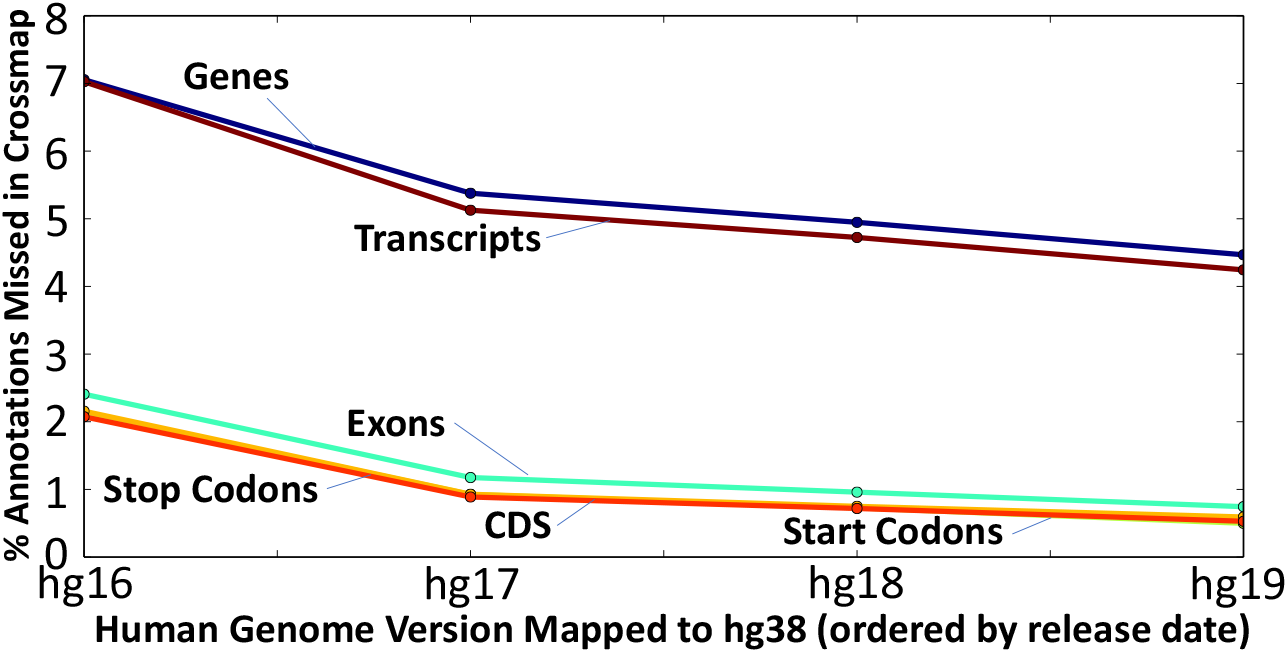
Percentage of different annotations missed when remapping reads from an old reference (x-axis) to the latest reference (hg38), using a remapping tool that solely relies on existing chain files (e.g., UCSC LiftOver [27]).

**Figure S2:**
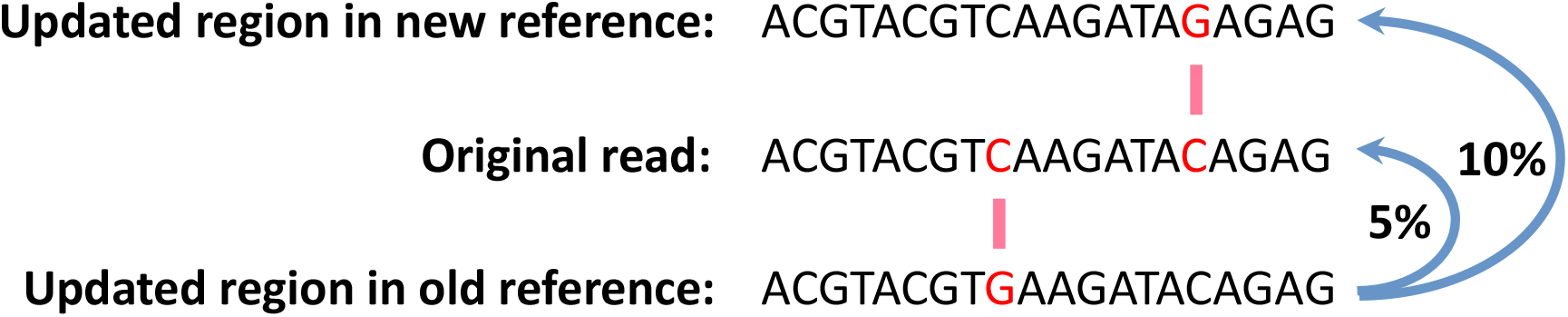
In order to comprehensively account for possible mappings of a read that previously mapped to an old reference genome, we create a lookup table describing the similarity between two reference genomes, using 2*×* the alignment error acceptance rate. As an example, if a read aligns to a location in the old reference genome with a 5% error rate (1 substitution in 20 base pairs), it is possible for the same read to map to a location in the new reference genome (with a 5% error rate) whose sequence is 10% different (2 substitutions in 20 base pairs) from the sequence in the old reference genome.

## Footnotes

1 These tools are described in more detail in Supplementary Section S1.

2 A BAM file is the binary version of a SAM file. A SAM file is a tabdelimited text file that contains sequence alignment data [37].

3 FastRemap implements necessary modifications to the *CrossMap* code such that its output is compatible with GATK (See Supplementary Section S5).

4 The GATK HaplotypeCaller tool cannot analyze the outputs of *CrossMap* or *LiftOver* since their outputs are not compatible with downstream analysis tools (as described in Supplementary Section S5). Therefore, we do not analyze the outputs of *CrossMap* or *LiftOver* in this section.

