## Supplemental Pages for "AirLift: A Fast and Comprehensive Technique for Remapping Alignments between Reference Genomes"

### Supplementary File for AirLift: A Fast and Comprehensive Technique for Translating Alignments between Reference Genomes

Jeremie S. Kim, Can Firtina, Meryem Banu Cavlak, Damla Senol Cali, Nastaran Hajinazar, Mohammed Alser, Can Alkan, and Onur Mutlu

### Supplementary Tables for AirLift

Table S1: Annotations in the new reference not covered by reads when remapping reads across reference genomes with UCSC LiftOver.

New Reference

**hg19**

| Old Ref. | gene | exon | stop codon | CDS | start codon | transcript |
|---|---|---|---|---|---|---|
| **hg16** | 3.07 | 0.92 | 0.79 | 0.77 | 0.72 | 2.92 |
| **hg17** | 1.45 | 0.36 | 0.23 | 0.24 | 0.24 | 1.22 |
| **hg18** | 0.84 | 0.12 | 0.07 | 0.10 | 0.10 | 0.78 |

**hg38**

| Old Ref. | gene | exon | stop codon | CDS | start codon | transcript |
|---|---|---|---|---|---|---|
| **hg16** | 7.06 | 2.41 | 2.13 | 2.16 | 2.07 | 7.03 |
| **hg17** | 5.38 | 1.18 | 0.93 | 0.93 | 0.89 | 5.13 |
| **hg18** | 4.95 | 0.96 | 0.73 | 0.75 | 0.72 | 4.72 |
| **hg19** | 4.47 | 0.74 | 0.50 | 0.59 | 0.53 | 4.24 |

Between each pair of reference genomes, we indicate the exact values of specific annotation types (e.g., gene, exon, stop codon, CDS, start codon, transcript) that are "lost" when using UCSC LiftOver [1] on a read data set from an old reference (rows) to a new reference (columns). Briefly, 3.07% of the gene model coordinates in hg16 assembly are not found in hg19, where the loss rate of genes reaches 4.47% between the most recent two assembly versions (hg19 and hg38).

Table S2: Details of the reference genomes that we use in our evaluations.

| Species | Version | Total # of Bases | non-N Bases | Release Date |
|---|---|---|---|---|
| Human | hg16 | 3,091,959,510 | 2,865,086,288 | 2004-02-04 |
| | hg17 | 3,091,360,260 | 2,865,812,574 | 2004-08-24 |
| | hg18 | 3,104,054,490 | 2,881,568,385 | 2006-03-03 |
| | hg19 | 3,137,144,693 | 2,897,293,955 | 2009-02-27 |
| | hg38 | 3,209,286,105 | 3,049,316,098 | 2013-12-24 |
| C. elegans | ce2 | 100,291,769 | 100,291,761 | 2004-03-01 |
| | ce4 | 100,281,244 | 100,281,244 | 2007-01-01 |
| | ce6 | 100,281,426 | 100,281,244 | 2008-05-01 |
| | ce10 | 100,286,070 | 100,286,070 | 2012-04-13 |
| | ce11 | 100,286,401 | 100,286,401 | 2013-02-07 |
| Yeast | sacCer1 | 12,156,302 | 12,156,302 | 2001-10-01 |
| | sacCer2 | 12,162,995 | 12,162,995 | 2008-06-01 |
| | sacCer3 | 12,157,105 | 12,157,105 | 2014-12-17 |

Table S3: Read data sets that we use in our evaluations.

| Read Data Set | Accession No. | Details |
|---|---|---|
| Human NA12878 - Illumina | ERR194147 | 795,505,905 paired-end reads (101bps each, 50× coverage) |
| Human NA12878 - Illumina | ERR262997 | 643,097,275 paired-end reads (101bps each, 40× coverage) |
| C. elegans N2 - Illumina | SRR3536210 | 78,696,056 paired-end reads (101bps each, 150× coverage) |
| Yeast S288C - Illumina | ERR1938683 | 3,318,467 paired-end reads (150bps each, 82× coverage) |

Table S4: AirLift Preprocessing Time

| Species | Remapping reads From | To | Preprocessing Time (s) | **Processing Time (s)** |
|---|---|---|---|---|
| Human | hg16 | hg38 | 10303.83 (1.88%) | **548075.93** |
| | hg17 | | 10005.79 (1.91%) | **523863.33** |
| | hg18 | | 7764.80 (1.61%) | **467759.21** |
| | hg19 | | 6912.65 (1.58%) | **437509.45** |
| C. elegans | ce2 | ce4 | 69.12 (1.33%) | **5197.01** |
| | ce4 | ce10 | 70.95 (1.32%) | **5374.81** |
| | ce6 | | 68.14 (1.29%) | **5281.88** |
| | ce6 | ce11 | 56.94 (1.04%) | **5475.44** |
| | ce10 | | 56.98 (1.07%) | **5324.87** |
| Yeast | sacCer1 | sacCer2 | 6.50 (2.43%) | **267.52** |
| | sacCer1 | sacCer3 | 10.22 (3.56%) | **287.08** |
| | sacCer2 | | 11.46 (4.23%) | **270.92** |

We show for our selected species' reference genomes, human (large), C. elegans (medium), yeast (small) the execution time for preprocessing a pair of references. The execution time is shown for different version pairs of each reference genome (row). The preprocessing time for each pair of references is measured in seconds and the percentage of the full remapping time (processing time) is shown in parenthesis.

Table S5: Breakdown of Region Labels for Each Pair of Reference Genomes.

| Species | Remapping a read set From | To | Constant (%) | Updated (%) | Retired (%) |
|---|---|---|---|---|---|
| Human | hg16 | hg38 | 85.7475 | 14.1867 | 0.0659 |
| | hg17 | | 86.5513 | 13.4106 | 0.0382 |
| | hg18 | | 86.6874 | 13.2485 | 0.0641 |
| | hg19 | | 87.1995 | 12.7344 | 0.0660 |
| C. elegans | ce2 | ce4 | 99.9862 | 0.0109 | 0.0028 |
| | ce4 | ce10 | 99.9738 | 0.0222 | 0.0040 |
| | ce6 | | 99.9770 | 0.0191 | 0.0040 |
| | ce6 | ce11 | 99.8262 | 0.1587 | 0.0151 |
| | ce10 | | 99.8505 | 0.1428 | 0.0067 |
| Yeast | sacCer1 | sacCer2 | 90.2503 | 8.7276 | 1.0220 |
| | sacCer1 | sacCer3 | 99.4449 | 0.5297 | 0.0254 |
| | sacCer2 | | 99.5459 | 0.4289 | 0.0252 |

We show for our selected species' reference genomes, human (large), C. elegans (medium), yeast (small) how versions of the reference genome (row) are comprised of distinct regions (i.e., constant, updated, retired) in relation to a more recent version of the species. Each cell contains the percentage of the old reference genome that each category of regions (columns) comprises.

Table S6: Execution Time Breakdown for Remapping a Read Set by Case

| Species | Remapping reads | | Time to remap reads that originally mapped to region (s): | | | | |
|---|---|---|---|---|---|---|---|
| | From | To | 1. Constant | 2. Updated | 3. Retired | 4. Unmapped | **Total** |
| Human | hg16 | hg38 | 107390.64 (19.59%) | 430851.66 (78.61%) | 98.59 (0.02%) | 9735.04 (1.78%) | **548075.93** |
| | hg17 | | 107373.59 (20.50%) | 408492.93 (77.98%) | 79.35 (0.02%) | 7917.46 (1.51%) | **523863.33** |
| | hg18 | | 106318.91 (22.73%) | 354026.40 (75.69%) | 77.36 (0.02%) | 7336.54 (1.57%) | **467759.21** |
| | hg19 | | 106136.64 (24.26%) | 324513.21 (74.17%) | 52.39 (0.01%) | 6807.21 (1.56%) | **437509.45** |
| C. elegans | ce2 | ce4 | 5138.04 (98.87%) | 8.23 (0.16%) | 0.09 (0.00%) | 50.65 (0.97%) | **5197.01** |
| | ce4 | ce10 | 5311.64 (98.82%) | 9.03 (0.15%) | 0.00 (0.00%) | 54.14 (1.01%) | **5374.81** |
| | ce6 | | 5219.78 (98.82%) | 7.96 (0.15%) | 0.00 (0.00%) | 54.14 (1.03%) | **5281.88** |
| | ce6 | ce11 | 5386.70 (98.38%) | 36.84 (0.67%) | 0.00 (0.00%) | 51.90 (0.95%) | **5475.44** |
| | ce10 | | 5273.38 (99.03%) | 0.00 (0.00%) | 0.00 (0.00%) | 51.49 (0.97%) | **5324.87** |
| Yeast | sacCer1 | sacCer2 | 263.90 (98.65%) | 2.60 (0.97%) | 0.00 (0.00%) | 1.42 (0.53%) | **267.52** |
| | sacCer1 | sacCer3 | 276.75 (96.40%) | 10.13 (3.53%) | 0.00 (0.00%) | 1.42 (0.50%) | **287.08** |
| | sacCer2 | | 262.32 (96.82%) | 8.13 (3.00%) | 0.00 (0.00%) | 1.42 (0.53%) | **270.92** |

We show for our selected species' reference genomes, human (large), C. elegans (medium), yeast (small) the execution time breakdown of remapping a read set. The execution time is shown for different versions of each reference genome (row) and is shown for the four remapping cases: 1) reads that originally mapped to a constant region, 2) reads that originally mapped to an updated region, 3) reads that originally mapped to a retired region, and 4) reads that never mapped to the old reference (i.e., unmapped). The remap time for each case is measured in seconds and the percentage of the full remapping time is shown in parenthesis.

Table S7: Number of Reads Remapped by Each AirLift Case

| Species | Remapping reads | | Time to remap reads that originally mapped to region (s): | | | | |
|---|---|---|---|---|---|---|---|
| | From | To | 1. Constant | 2. Updated | 3. Retired | 4. Unmapped | **Total** |
| Human | hg16 | hg38 | 2866050090 (86.57%) | 401163904 (12.12%) | 408920 (0.0124%) | 43205312 (1.30%) | **3310828226** |
| | hg17 | | 2869062015 (86.78%) | 396473981 (11.99%) | 126607 (0.0038%) | 40431845 (1.22%) | **3306094448** |
| | hg18 | | 2870008151 (88.04%) | 350216312 (10.74%) | 126040 (0.0039%) | 39551799 (1.21%) | **3259902302** |
| | hg19 | | 2870919584 (90.88%) | 249423596 (7.90%) | 60300 (0.0019%) | 38781755 (1.23%) | **3159185235** |
| C. elegans | ce2 | ce4 | 155088879 (98.30%) | 59432 (0.038%) | 1700 (0.002%) | 2413178 (1.53%) | **157563189** |
| | ce4 | ce10 | 155091089 (98.41%) | 91980 (0.058%) | 0 (0.000%) | 2413157 (1.53%) | **157596226** |
| | ce6 | | 155091089 (98.46%) | 9444 (0.006%) | 0 (0.000%) | 2413157 (1.03%) | **157513690** |
| | ce6 | ce11 | 155091090 (98.47%) | 1040 (0.001%) | 0 (0.000%) | 2413156 (1.53%) | **157505286** |
| | ce10 | | 155094686 (98.47%) | 0 (0.000%) | 0 (0.000%) | 2409560 (1.53%) | **157504246** |
| Yeast | sacCer1 | sacCer2 | 6509230 (97.79%) | 17676 (0.27%) | 0 (0.00%) | 129581 (1.95%) | **6656487** |
| | sacCer1 | sacCer3 | 6509230 (97.36%) | 63404 (0.95%) | 0 (0.00%) | 129581 (1.93%) | **6702215** |
| | sacCer2 | | 6509230 (97.79%) | 46652 (0.70%) | 0 (0.00%) | 129576 (1.94%) | **6685458** |

We show for our selected species' reference genomes, human (large), C. elegans (medium), yeast (small) the number of reads in a read set that is mapped by each AirLift case. The number of reads is shown for different versions of each reference genome (row) and is shown for the four remapping cases: 1) reads that originally mapped to a constant region, 2) reads that originally mapped to an updated region, 3) reads that originally mapped to a retired region, and 4) reads that never mapped to the old reference (i.e., unmapped). The percentage of the full read set is shown in parenthesis.

Table S8: Execution Time per Read when Remapping a Read Set by Case

| Species | Remapping reads | | Avg time to remap a read originally mapped to region (us) | | | |
|---|---|---|---|---|---|---|
| | From | To | 1. Constant | 2. Updated | 3. Retired | 4. Unmapped |
| Human | hg16 | hg38 | 37.470 | 1074.004 | 241.099 | 225.320 |
| | hg17 | | 37.425 | 1030.315 | 626.743 | 195.822 |
| | hg18 | | 37.045 | 1010.879 | 613.773 | 185.492 |
| | hg19 | | 36.970 | 1301.053 | 868.822 | 175.526 |
| C. elegans | ce2 | ce4 | 33.130 | 138.478 | 52.941 | 20.989 |
| | ce4 | ce10 | 34.249 | 98.174 | - | 22.435 |
| | ce6 | | 33.656 | 842.863 | - | 22.435 |
| | ce6 | ce11 | 34.732 | 3542.308 | - | 21.507 |
| | ce10 | | 33.812 | - | - | 21.369 |
| Yeast | sacCer1 | sacCer2 | 40.542 | 14.709 | - | 10.958 |
| | sacCer1 | sacCer3 | 42.517 | 15.977 | - | 10.958 |
| | sacCer2 | | 40.300 | 17.427 | - | 10.959 |

We show for our selected species' reference genomes, human (large), C. elegans (medium), yeast (small) the average execution time to remap a single read depending on the AirLift case. The execution time is shown for different versions of each reference genome (row) and is shown for the four remapping cases: 1) reads that originally mapped to a constant region, 2) reads that originally mapped to an updated region, 3) reads that originally mapped to a retired region, and 4) reads that never mapped to the old reference (i.e., unmapped). The average remap time for each read is measured in microseconds. Each value is calculated by dividing the corresponding cell in Table S6 by the corresponding cell in Table S7.

### Supplementary Figures for AirLift

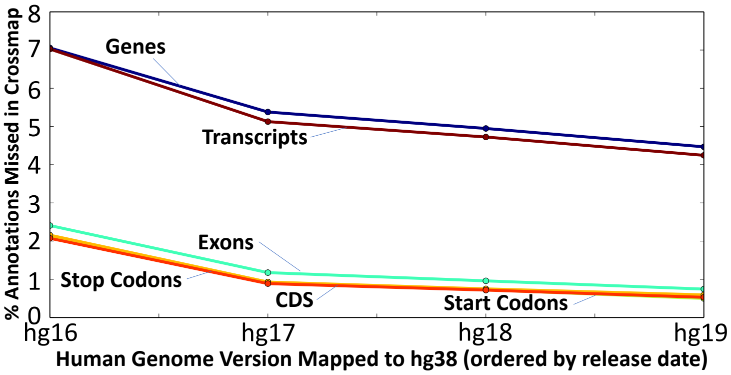


Figure S1: Percentage of different annotations missed when remapping reads from an old reference (x-axis) to the latest reference (hg38), using UCSC LiftOver [1].

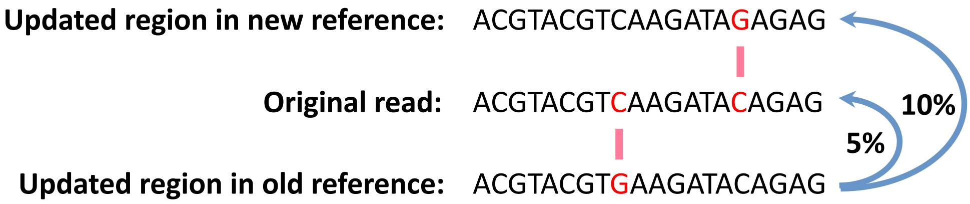


Figure S2: In order to comprehensively account for possible mappings of a read that previously mapped to an old reference genome, we create a lookup table describing the similarity between two reference genomes, using 2× the alignment error acceptance rate. As an example, if a read aligns to a location in the old reference genome with a 5% error rate (1 substitution in 20 base pairs), it is possible for the same read to map to a location in the new reference genome (with a 5% error rate) whose sequence is 10% different (2 substitutions in 20 base pairs) from the sequence in the old reference genome.

### References

[1] UCSC. UCSC LiftOver: Lift Genome Annotations. https://genome.ucsc.edu/cgi-bin/hgLiftOver.

[2] UCSC. UCSC Genome Browser: Sequence and Annotation Downloads. http://hgdownload.soe.ucsc.edu/downloads.html.

[3] Hao Zhao, Zhifu Sun, Jing Wang, Haojie Huang, Jean-Pierre Kocher, and Liguo Wang. CrossMap: A Versatile Tool for Coordinate Conversion Between Genome Assemblies. *Bioinformatics*, 30(7):1006–1007, 2013.

[4] Zhao, Hao and Sun, Zhifu and Wang, Jing and Huang, Haojie and Kocher, Jean-Pierre and Wang, Liguo. CrossMap: Convert Genome Coordinates Between Assemblies. http://crossmap.sourceforge.net/#use-pip-to-install-crossmap.

[5] NCBI. NCBI Genome Remapping Service. https://www.ncbi.nlm.nih.gov/genome/tools/remap.

[6] Bo Gao, Qingyao Huang, and Michael Baudis. Segment_Liftover: A Python Tool to Convert Segments Between Genome Assemblies. *F1000Research*, 7, 2018.

[7] Bo Gao. Segment Liftover. https://pypi.org/project/segment-liftover/.

[8] Belinda Giardine, Cathy Riemer, Ross C Hardison, Richard Burhans, Laura Elnitski, Prachi Shah, Yi Zhang, Daniel Blankenberg, Istvan Albert, James Taylor, et al. Galaxy: A Platform for Interactive Large-scale Genome Analysis. *Genome Research*, 15(10):1451–1455, 2005.

[9] The Galaxy Team. Galaxy. https://www.usegalaxy.org.

[10] Konstantin Tretyakov. Pyliftover. https://pypi.org/project/pyliftover/.

[11] Simon P Sadedin and Alicia Oshlack. Bazam: A Rapid Method for Read Extraction and Realignment of High-Throughput Sequencing Data. *Genome Biology*, 20(1):78, 2019.

[12] Heng Li and Richard Durbin. Fast and Accurate Short Read Alignment with Burrows–Wheeler Transform. *Bioinformatics*, 25(14):1754–1760, 2009.

[13] Chain Format. https://genome.ucsc.edu/goldenPath/help/chain.html.

[14] Xiangqun Zheng-Bradley, Ian Streeter, Susan Fairley, David Richardson, Laura Clarke, Paul Flicek, and 1000 Genomes Project Consortium. Alignment of 1000 Genomes Project Reads to Reference Assembly GRCh38. *GigaScience*, 6(7):1–8, 2017.
